# Integration of taxonomic signals from MAGs and contigs improves read annotation and taxonomic profiling of metagenomes

**DOI:** 10.1101/2023.03.22.533753

**Authors:** Ernestina Hauptfeld, Nikolaos Pappas, Sandra van Iwaarden, Basten L. Snoek, Andrea Aldas-Vargas, Bas E. Dutilh, F.A. Bastiaan von Meijenfeldt

## Abstract

Metagenomic analysis typically includes read-based taxonomic profiling, assembly, and binning of metagenome-assembled genomes (MAGs). Here we integrate these steps in Read Annotation Tool (RAT), which uses robust taxonomic signals from MAGs and contigs to enhance read annotation. RAT reconstructs taxonomic profiles with high precision and sensitivity, outperforming other state-of-the-art tools. In high-diversity groundwater samples, RAT annotates a large fraction of the metagenomic reads, calling novel taxa at the appropriate, sometimes high taxonomic ranks. Thus, RAT integrative profiling provides an accurate and comprehensive view of the microbiome. CAT/BAT/RAT is available at https://github.com/MGXlab/CAT. The CAT pack now also supports GTDB annotations.

## Background

Metagenomic shotgun sequencing provides a single platform for exploring both the composition and the functional potential of diverse microbial communities [1–5]. While functional profiling maximizes the usage of the shotgun data, taxonomic profiling of metagenomes may involve mapping reads to a reference database containing specific marker genes [6–10], in which case only a portion of the data is used and function can only be coupled to taxonomy for those reads that contain the marker gene. Alternatively, taxonomy can be assigned to as much of the data as possible by querying reads to a full reference database [11–14]. Metagenomic profilers carry out direct homology searches in DNA [12], protein [13], or k-mer space [11,15,16], and the resulting taxonomic profiles have been used in large scale studies to characterize microbial communities of the oceans [1], the global topsoil [17], and to estimate the niche range of every known microbial taxon [18].

Taxonomic profiles that are based on direct queries of individual reads to full reference databases give a comprehensive view of a microbiome, but often contain spurious annotations. Assigning taxonomy based on homology searches is challenging, particularly for relatively short reads: (i) some genomic regions are highly conserved across taxa, making it difficult to discriminate between them; (ii) microbes have high rates of horizontal gene transfer [19,20], so the best hit in the reference database might be an unrelated taxon; (iii) environmental microbiomes may contain many novel taxa without close representatives in the reference database, resulting in possible annotation to e.g. a genus or species when the organism only shares the same order [21,22]; (iv) known taxa may contain novel genomic regions, resulting in no annotation of reads covering that region or annotation to a more distant relative, and (v) reference databases contain mis-annotated sequences [23]. These challenges are especially pronounced when directly comparing individual reads. With the exception of data from recent long-read sequencing platforms [24], reads are short sequences that contain limited taxonomic information, leading to reads derived from a single strain potentially being assigned to several different taxa. Thus, while comprehensive, taxonomic profiles based on read annotations are inherently noisy with spurious annotations, and often inaccurate [21].

Over the past decade, best practices in shotgun metagenomics have been established, including reference database-independent (*de novo*) assembly [25,26] and binning of metagenome assembled genomes (MAGs) [27,28]. The resulting contiguous sequences (contigs) and especially MAGs allow for accurate detection of novel taxa. Contigs and MAGs are significantly longer than the original short sequencing reads, the additional data allowing for more reliable taxonomic annotation, either by multiple homology searches [29,30] or phylogenetic placement [31]. Long sequence length mitigates the errors in annotation discussed earlier because multiple taxonomic signals can be integrated (confidence in annotation: MAGs > contigs > reads). However, even though taxonomic annotation is more accurate for longer sequences, they often represent only part of the metagenomic data and therefore provide an incomplete picture of the microbiome (data explained: reads > contigs > MAGs). As *de novo* assembly and binning depends on sufficient coverage of the genome sequence, it may be expected that especially rare microorganisms will be missed when MAGs or contigs are assessed. For a robust taxonomic profile that also includes rare microorganisms, an annotation protocol that integrates both taxonomic information from long sequences where available and short reads where not may thus be desirable.

Here, we present Read Annotation Tool (RAT), an annotation pipeline for metagenomic sequencing reads that integrates accurate annotation of contigs and MAGs derived from *de novo* assembly and binning, and direct homology searches of the remaining unassembled reads. RAT estimates taxonomic profiles by associating reads to longer sequences when possible and assigning taxonomy according to the most reliable taxonomic signal it can find (MAGs > contigs > reads). Contigs and MAGs are taxonomically annotated with the previously published tools CAT and BAT [29], which provide robust annotation based on open reading frame (ORF) prediction and comparisons to a protein database [32–34]. We show that, by integrating taxonomic signals from MAGs, contigs, and reads, RAT provides more accurate read annotations and taxonomic profiles than other state-of-the-art tools, and accurately characterizes groundwater microbiomes with many novel taxa.

## Results and discussion

Natural microbial communities consist of many different microorganisms that can be identified and characterized by sequencing their DNA with shotgun metagenomics. To get an accurate overview of all microorganisms and their relative abundances in a sample, the most comprehensive approach is to obtain reliable taxonomic annotation for as many of the sequencing reads as possible. While contigs and MAGs can be more reliably annotated than individual reads, in most metagenomic datasets not all reads are assembled into contigs and not all contigs are binned into MAGs (**Figure 1**). To address this trade-off between annotation accuracy and the fraction of data that can be explained in a metagenome, we developed Read Annotation Tool (RAT) (**Figure 1B,C**).

**Figure 1.**
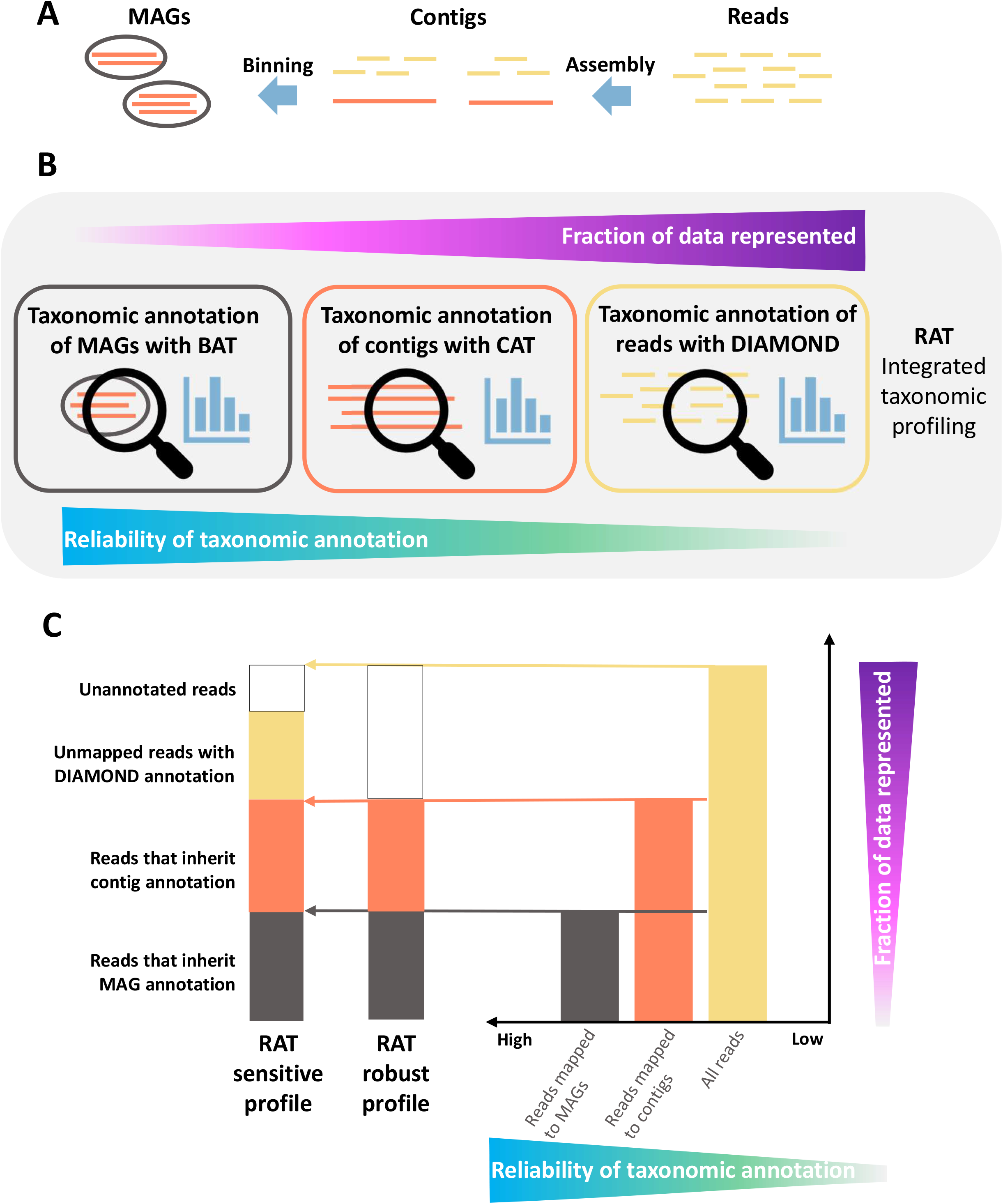
The RAT workflow. (A) Overview of a standard state-of-the-art metagenomics pipeline. (B) Overview of the RAT workflow: Reads are mapped to contigs which are binned into MAGs or unbinned. MAGs and contigs are taxonomically annotated using BAT/CAT. Unmapped reads and unclassified contigs are annotated using DIAMOND. (C) Left: Composition of an integrated taxonomic profile as reconstructed by RAT sensitive and RAT robust. Right: Schematic bar plot showing the fraction of the metagenome that can be annotated as reads, contigs, and MAGs.

RAT annotates contigs and MAGs with the previously published tools Contig Annotation Tool (CAT) and Bin Annotation Tool (BAT), respectively. CAT and BAT query predicted ORFs on these longer sequences to a protein reference database with DIAMOND blastp [33], and assign taxonomy based on the combined taxonomic signal [22]. Default options for the reference database include the non-redundant protein database (nr) [35] and in the latest update the non-redundant set of proteins in the Genome Taxonomy Database (GTDB) [36], and alternatively any protein database with taxonomic annotations can be supplied by the user. Next, individual reads are mapped to the contigs and each read inherits the taxonomic annotation with the highest confidence. Finally, the remaining sequences (reads that do not map to a contig and contigs that cannot be annotated by CAT) are annotated individually by querying them to the protein database with DIAMOND blastx [33]. Thus, by assigning reads to taxonomic annotation with the highest confidence, RAT reconstructs a comprehensive taxonomic profile with high accuracy (**Figure 1C**). The final step in which sequences are individually queried to the protein database is optional, and depending on whether this step is included, we distinguish two RAT modes: in ‘robust’ mode, RAT only uses the most reliable read annotations, which are based on MAGs and contigs with ORFs. In ‘sensitive’ mode, RAT also uses the read and contig annotations with DIAMOND blastx, which also include more tentative annotations while representing more of the data.

We evaluated the performance of RAT for read annotation, and how well the final taxonomic profile represents the microbial community. First, to address the trade-off between the annotation accuracy and the fraction of reads that can be annotated by the different steps in RAT, we used simulated data from the Critical Assessment of Metagenome Interpretation (CAMI) challenge [37]. Second, we used the same dataset to compare taxonomic profiles predicted by RAT to those predicted by other commonly used state-of-the-art profilers. Third, we assessed the performance of RAT and the other profilers on real metagenomes. To this end, we analyzed samples from three groundwater monitoring wells, a relatively unexplored high-diversity environment that contains many novel taxa [38].

### Including taxonomic signals from MAGs and contigs improves read annotation

To evaluate how the integration of different taxonomic signals influences the annotation of individual reads, we annotated simulated metagenomic datasets from the second CAMI challenge [37] with RAT. The CAMI challenge simulated well-characterized microbiomes of the mouse gut. The 10 datasets contained between 97-225 species, and included raw reads, gold standard assemblies (the best possible assembly of the sequencing reads in a sample), and genome sequences of these species. In our benchmarks, we used the gold standard assemblies as contig input.

We compared five different methods for read annotation: (i) we annotated all reads directly with DIAMOND blastx, without mapping them to contigs or MAGs, (ii) We ran RAT sensitive using only contig annotations, and direct read annotations for reads that do not map to contigs, (iii) We ran RAT sensitive also using MAG annotations with the genome sequences included in the dataset (‘CAMI genomes’) as input, (iv) We ran RAT sensitive with MAG annotations using MAGs binned by MetaBAT2 [27], and (v) we ran RAT robust using only contig and MetaBAT2 MAG annotations but no direct read annotations (**Figure 2**). Results were assessed at six taxonomic ranks (phylum, class, order, family, genus, and species) and we scored whether a read was correctly or incorrectly annotated, or unclassified (**Figure 2**).

**Figure 2.**
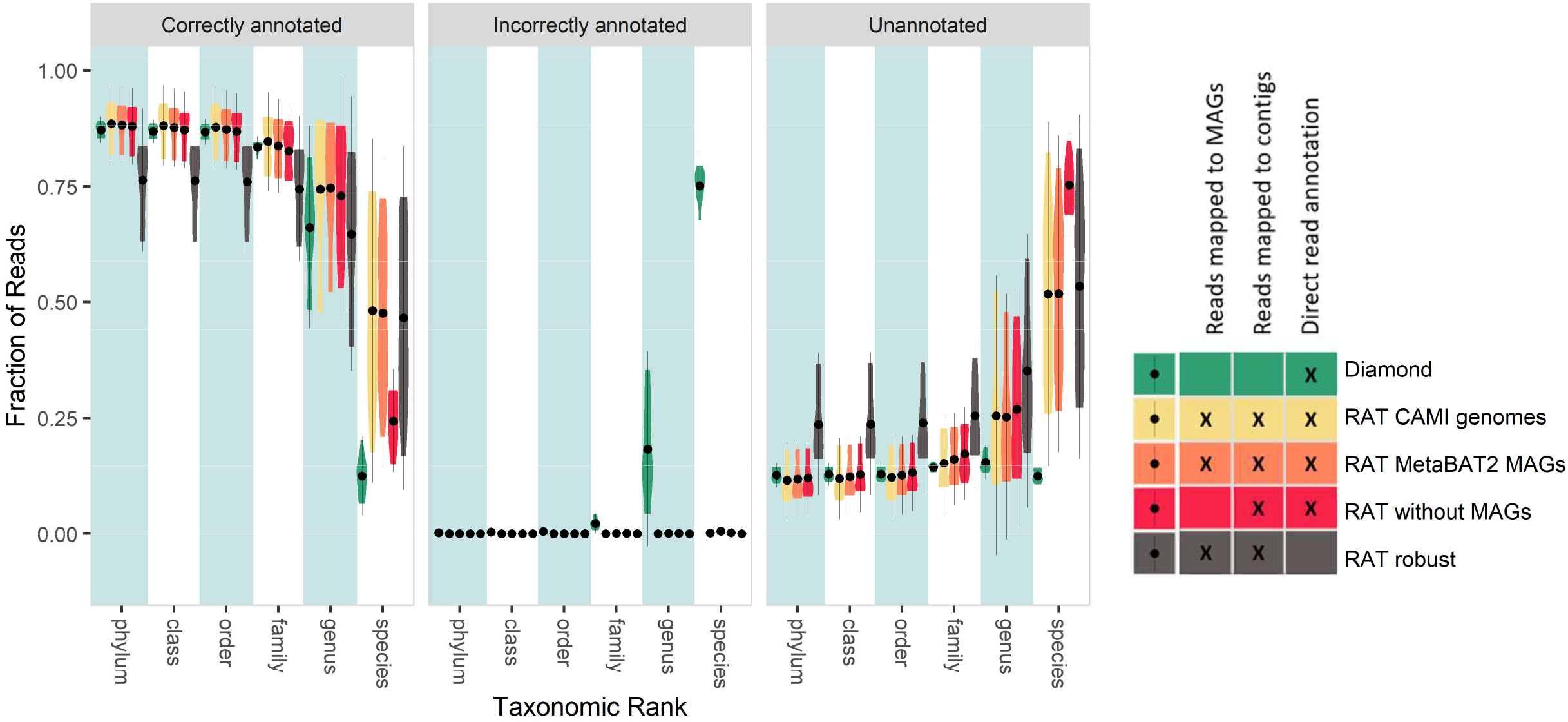
Outcome of incorporating different taxonomic signals into read annotations. ‘Diamond’ refers to using only direct read annotation. ‘RAT CAMI genomes’ refers to a RAT sensitive run using the genomes that were provided by the CAMI challenge as MAG input. ‘RAT MetaBAT2 MAGs’ refers to a RAT sensitive run with contigs binned by MetaBAT2. ‘RAT without MAGs’ refers to a RAT sensitive run without MAG input. ‘RAT robust’ refers to a RAT robust run, using only read annotation via mapping to MetaBAT2 MAGs and contigs, but no direct read annotation. In the left panel, a higher value is better, in the middle and right panel, a lower value is better.

Direct annotation with DIAMOND blastx resulted in low accuracy at low taxonomic ranks with a high fraction of mis-annotated reads (**Figure 2**), revealing spurious annotations when mapping short sequences to a reference database. Accuracy is particularly low on species rank, where only 12.4±4% (mean ± standard deviation) of the reads were correctly annotated by DIAMOND. Despite using DIAMOND with the same reference database, RAT runs reduced mis-annotations and improved the fraction of correctly annotated reads at all taxonomic ranks, highlighting the value of integrating information from taxonomically annotated MAGs and contigs (**Figure 2**).

When only taxonomic signals from contigs are integrated, the fraction of correctly annotated and unclassified reads increases compared to direct annotation with DIAMOND blastx, while the fraction of incorrectly annotated reads drops to 0.1-1%. This indicates that many previously mis- or unannotated reads are correctly annotated if they map to contigs. The fact that many of the reads that were mis-annotated using DIAMOND blastx are unclassified in RAT sensitive shows that these reads are mapped to contigs that cannot be annotated on lower taxonomic ranks by CAT. This indicates that the contigs have hits to multiple different taxa in the database, in which case CAT chooses a higher rank taxon as the most robust taxonomic annotation [22], precluding annotation on lower taxonomic ranks.

When taxonomic signals from both contigs and MAGs are integrated, the fraction of correctly annotated reads increases while the fraction of unclassified reads decreases compared to annotating without MAGs. In the CAMI mouse gut dataset, using the CAMI genomes as MAG input and binning the contigs with MetaBAT2 gave very similar results, indicating that current binning tools accurately group contigs from the same species together. Without using DIAMOND blastx to annotate the remaining unmapped reads and unclassified contigs (RAT robust), the fraction of annotated reads drops, while the fraction of unclassified reads increases. This effect is likely more pronounced in some real biological datasets, where higher taxon diversity makes it more difficult to assemble reads into contigs than in the simulated CAMI samples (for which a gold standard assembly is supplied), and which thus contain more unmapped reads (see **below**, **Supplementary Figures S3 & S4**).

Concluding, using the taxonomic signals from contigs and MAGs for read annotation leads to more reliable annotations than using direct querying of individual reads.

### Including information from contigs and MAGs improves accuracy of taxonomic profiling

Metagenomics is used to analyze high complexity microbial communities, including many different taxa with orders of magnitude of difference in their abundances. Taxonomic profilers aim to chart the community composition by listing all taxa in a sample and estimate their relative abundance. A good taxonomic profile contains as many members of the microbial community as possible, while avoiding taxa that are not present in the sample. In practice, this often leads to a compromise between sensitivity (finding all taxa that are present and maybe some false positives) and precision (avoiding taxa that are not present and maybe some false negatives). To assess how including contig and MAG annotations affects the reconstruction of taxonomic profiles, we used four metrics (sensitivity, precision, L1 distance, and weighted UniFrac distance) to compare the CAMI ground truth taxonomic profiles to those reconstructed by RAT and four state-of-the-art taxonomic profilers that carry out annotations by direct read mapping (**Figure 3**). Centrifuge classifies microbial DNA sequences, Kaiju annotates sequences in protein space, Kraken2 annotates DNA sequences using exact kmer matches, and Bracken uses Kraken2 annotations for a bayesian reestimation of the abundances of taxa in the sample. As direct read annotations can be inaccurate, we limited the amount of noise by only counting taxa that were detected in the profile with a minimum relative abundance of 0.001%.

**Figure 3.**
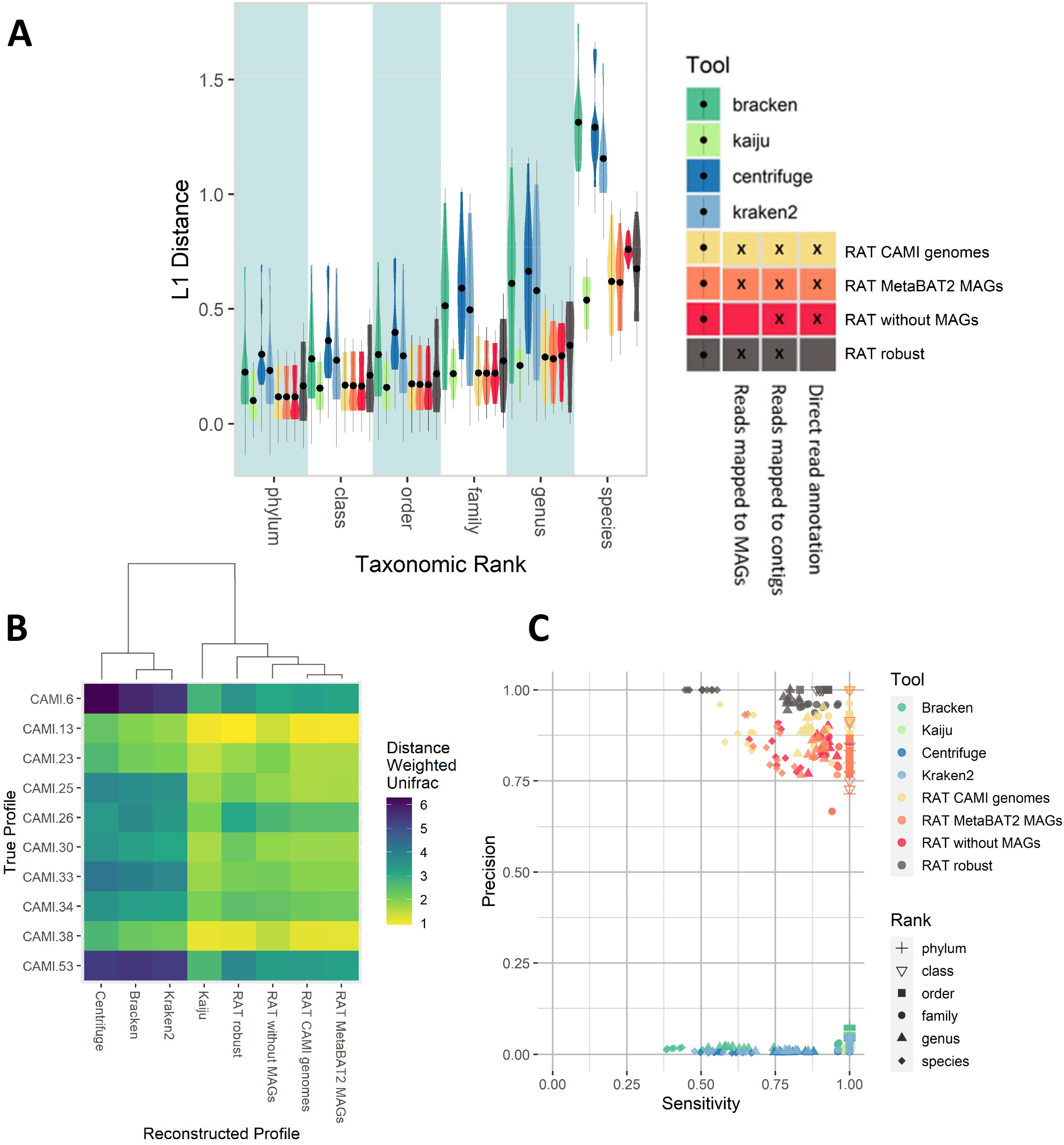
Similarities between true profiles and profiles reconstructed by different tools of the CAMI2 mouse gut dataset. We only counted taxa as detected if their relative abundance was at least 0.001%. (A) L1 Values between profiles reconstructed by RAT/other tools and the true profiles. An L1 value of 0 means that two profiles are identical (thus lower is better). (B) Heatmap of weighted Unifrac distances between reconstructed and true profiles (a shorter distance is better). (C) Sensitivity vs. precision of the different tools. Different shapes signify the sensitivity/precision on different taxonomic ranks, different colors indicate tools (high precision + sensitivity is better).

In line with our first benchmark, incorporating taxonomic signals from MAGs led to more accurate profiles than using only taxonomic signals from contigs, as seen in the L1 distance and in the weighted Unifrac distance. RAT sensitive slightly outperformed RAT robust (**Figure 3**) in L1 distance and sensitivity, indicating that including direct read annotation leads to reconstructed profiles that are more similar to the ground truth profile than when the step is not included. Taxonomic profiles reconstructed by RAT consistently had lower L1 distances to the ground truth profiles than profiles reconstructed by Bracken, Centrifuge, and Kraken2 (**Figure 3A**). In comparison to taxonomic profiles reconstructed by Kaiju, RAT runs had slightly higher L1 distances on genus and species rank. Taxonomic profiles reconstructed by RAT had lower weighted Unifrac distances to the ground truth profiles than Bracken, Centrifuge, and Kraken2 (**Figure 3B**) while Kaiju performed similarly.

RAT had a higher precision on all taxonomic ranks than the other evaluated tools. This means that RAT had fewer falsely detected taxa, in line with earlier observations of high precision of CAT and BAT annotations [22]. RAT robust maintained >0.94 precision on all taxonomic ranks, even when detected taxa were not limited by a minimum relative abundance cut-off (**Supplementary Figure S2**). Thus, like CAT and BAT on which its annotations are based, RAT robust tends to avoid spurious annotations at deeper taxonomic ranks in cases where conflicting taxonomic signals arise. For RAT sensitive, precision remained higher than that of the other evaluated tools across taxonomic ranks, albeit was lower than that of RAT robust. The minimum relative abundance cut-off greatly improved precision of RAT sensitive (cf. **Figure 3C** and **Supplementary Figure S2**). Spurious annotations are introduced wherever short sequencing reads are directly annotated, in the direct annotation step of RAT and in the other evaluated tools. However, because of the prioritization of taxonomic signals in RAT, a smaller fraction of total reads is annotated directly, leading to fewer spurious annotations in the first place. By setting an abundance cut-off (e.g. 0.001% of reads as in this benchmark), RAT can profit from the high sensitivity of the DIAMOND blastx step (finding taxa that might not be detected using just contig and MAG annotations) while further minimizing the number of falsely detected taxa (by excluding spurious annotations that have very low abundance).

RAT’s overall high precision can be explained by the integrated taxonomic profiling approach, which improves annotations in most of the challenges discussed above. Reads that map to conserved or horizontally transferred regions, or map to novel genomic regions of a known taxon are likely to get the correct annotation with RAT, because the surrounding (regions of the) genome is considered in the annotation via the contig and/or MAG. Reads belonging to novel taxa within known clades are also more likely to get correctly annotated, as when the reads are assembled into contigs or MAGs, RAT may annotate on a higher, appropriate taxonomic rank instead of on the lower taxonomic rank of closely related taxa. The difference in precision between the different approaches shows that reads that are not annotated by being associated with a contig or MAG, are far more likely to get falsely annotated. RAT’s approach reduces the number of falsely detected taxa from 200-4000 by the other evaluated tools to between 0 (RAT robust) and 38 (RAT sensitive).

All evaluated tools showed high sensitivity from phylum down to family rank, detecting most of the taxa that were present in the ground truth profiles (Figure 3C). This is consistent with increased barriers to horizontal gene transfer at higher taxonomic ranks [39]. On the genus and species ranks, RAT and Kaiju outperformed Bracken, Centrifuge, and Kraken2, while Kaiju showed higher sensitivity. RAT’s high performance on the CAMI datasets is in part due to the fact that a large fraction of the reads map back to contigs (81.6±6.6%) and MAGs (75.8±6.4%).These numbers are often lower in metagenomic datasets from other environments (see below). This leads to most reads being annotated in the most reliable MAG and contig annotation steps and few reads being annotated directly with DIAMOND, reducing the probability of spurious annotations (Supplementary Figures S3 & S4).

### Usage, runtime and memory requirements

Next, we compared the runtime and memory requirement of RAT with the other tools (**Table 1**). We did not take assembly and binning steps into account, since RAT does not assemble and bin metagenomes but rather takes assembled contigs and associated MAGs (of advised medium to high quality, see **below**) as input from the user. Although assembly and contig binning can take hours or days to run, they are a common procedure in many metagenomics studies, as they provide valuable genomic context information to short sequencing reads with relatively little risk of generating chimeras [40].

**Table 1.**
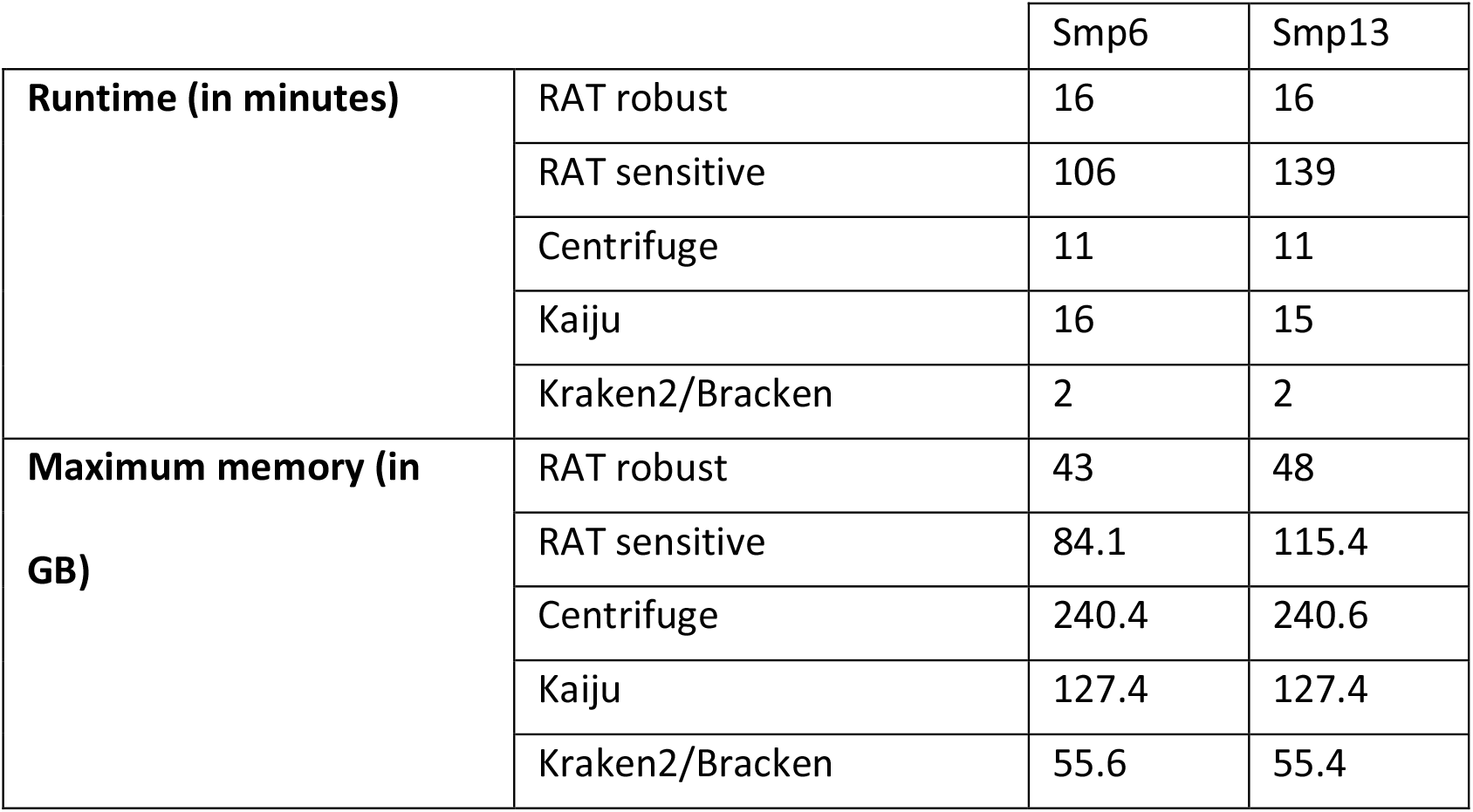
Runtime and Memory usage for RAT and four other tools. Tools were run on two simulated datasets (smp6: 33,098,456 reads, smp13: 33,184,772 reads) from the CAMI 2 challenge. RAT in robust mode (annotation based only on contigs and MAGs), and RAT in sensitive mode (annotation also based on DIAMOND annotation of unmapped reads). Kraken2 and bracken are run together. All runs were performed with 16 parallel CPUs.

Kraken2 was the fastest tool (01m49s), RAT sensitive was the slowest (02h05m10s), and all other tools including RAT robust performed the jobs in 16 minutes or less. In terms of memory usage, all tools can be run on a 256Gb server. RAT sensitive had a higher memory footprint than Kraken2, but lower than Kaiju and Centrifuge. RAT sensitive varied in RAM and runtime between the two samples because it loads different amounts of unclassified reads and contigs into memory depending on the sample.

### The expanded CAT pack facilitates the detection and annotation of unknown microorganisms

The simulated data provided by the CAMI challenge differs from real biological datasets. In the CAMI data, Illumina sequencing experiments were simulated of relatively low-diverse microbiomes containing genomes of known species. Annotations are facilitated by the fact that on average >80% of the reads mapped back to a MAG or contig from a gold-standard assembly, while in biological datasets, this percentage can be much lower (**Supplementary Figures S3 & S4**). In addition, particularly in microbiomes from under-studied environments, unknown lineages are often detected that are only distantly related to known taxa in reference databases. Awaiting taxonomic classification of these microorganisms, a higher-rank taxonomic annotation of the sequence at e.g. family or phylum rank may be appropriate in these cases.

RAT provides a framework for assessing these “unknowns”. Because reads are classified via CAT and BAT, annotations are made at the appropriate taxonomic rank. CAT and BAT assign individual ORFs to the last common ancestor of all hits that have a similar bit-score to the best hit, and annotate the contig or MAG using a bit-score based voting scheme that selects the taxon at which a certain fraction (in the RAT workflow the majority) of the ORF assignments agree [22]. Novel sequences have many distinct hits and are thus only annotated at a high taxonomic rank, reflecting their unknownness. MAGs that only receive a high taxonomic rank annotation by BAT may be further investigated with phylogenomic software for strain-level resolution [41,42]. Since the quality of RAT results is highly dependent on the quality of the input data, we recommend using high-quality assemblies, and only including MAGs with low contamination (e.g. <10% contamination according to CheckM [43]). Contaminated MAGs can be mis-annotated or annotated at a high trivial taxonomic rank in which case a contig annotation is more reliable. MAG completeness is less relevant for RAT, as MAGs with low completeness typically still include more than one contig from the same microorganism, creating a stronger taxonomic signal than present on the individual contigs.

To challenge RAT with real datasets, we selected relatively unexplored groundwater samples taken 12-64m below surface level from three different monitoring wells in a Dutch agricultural area, which we previously found had high microbial diversity and contain many novel taxa [38]. We performed a metagenomic analysis including quality control [44] assembly [25], and binning [27,28,45], which produced 514 MAGs. We supplied the reads, 2,770,251 contigs, and 423 medium- to high-quality MAGs (>50% completeness, <10% contamination [46]) to RAT to reconstruct taxonomic profiles of the groundwater samples, using nr as a reference database. In addition, the medium- to -high quality MAGs were dereplicated [47], and the resulting 195 representative MAGs were placed in a phylogenetic tree showing their relationships and abundance across samples (**Supplementary Figure S5**).

RAT annotated 22.0±8.7% (mean ± standard deviation) of reads by mapping them to MAGs, much less than in the simulated CAMI mouse gut datasets (see **Supplementary Figures S3 & S4**), reflecting the high complexity of the groundwater samples. RAT classified 20.9±3.2% of reads via unbinned contigs annotated by CAT, and 0.35±0.23% via contigs annotated by DIAMOND. Finally, DIAMOND blastx annotated an additional 23.0±3.3% of the reads. These unmapped reads represent sequences with low coverage that could not be assembled into contigs, and based on the results with simulated data above, we expect represent more spurious results. The taxonomic profile reconstructed by RAT sensitive showed that most reads belonged to unclassified Bacteria, including of the phyla *Chloroflexi* and *Deltaproteobacteria* (**Figure 4A**). *Chloroflexi* bacteria utilize a variety of electron acceptors including oxidized nitrogen or sulfur compounds. Comparison of the 18 reconstructed taxonomic profiles showed that Sample 23-2 contained relatively many *Chloroflexi* reads, while the *Deltaproteobacteria* were rare. Although many of the microorganisms in this sample could only be classified on high taxonomic ranks, 22 MAGs from these phyla represented 31.1% of the reads in the sample (see Supplementary Figure S5).

**Figure 4.**
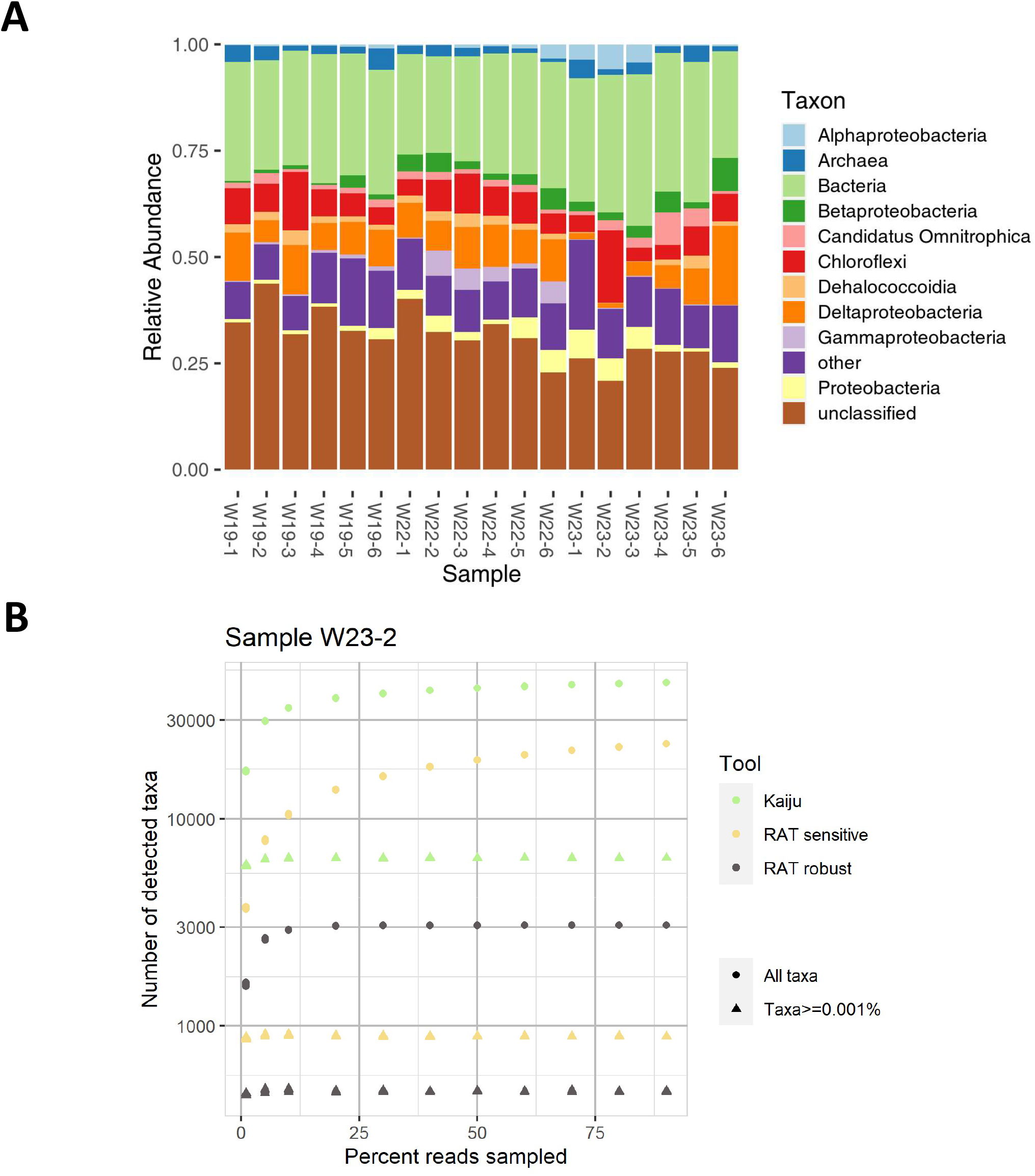
Taxonomic profiling of groundwater metagenomes. (A) Microbial profiles of groundwater samples on taxonomic rank class as reconstructed by RAT sensitive. (B) Rarefaction curves of the number of taxa detected in sample W23-2 by RAT robust, RAT sensitive, and Kaiju. Triangles indicate the number of taxa detected in profiles when a minimum abundance is required to consider an organism as detected. Circles indicate the number of taxa detected without a cut-off.

Next, we compared the taxonomic profiles of the groundwater metagenomes as predicted by RAT and Kaiju, as it was the best performing other tool in the previous benchmark. Both tools classified two thirds of the data (RAT: 68.9±5.8% of reads, Kaiju: 63.8±5.6% of reads, **Supplementary Table ST2**). However, RAT classified these reads into roughly 20% of the taxa that Kaiju predicted (**Figure 4B**). Bearing in mind the high precision of RAT (**Figure 3**), we propose that the taxa predicted by RAT are a more parsimonious interpretation of the metagenomic data than those predicted by Kaiju. To visualize the potential overestimation of taxa due to spurious annotations, we made rarefaction curves for RAT sensitive, RAT robust, and Kaiju. Without a minimum relative abundance cut-off, rarefaction curves for RAT sensitive and Kaiju did not level off. This reflects the spurious annotations of individual reads and indicates that, without a cut-off, deeper sequencing of the same sample would lead to higher predicted richness. This pattern was also observed in simulated data containing a known number of 110 species (**Supplementary Figure S6**). The rarefaction curve of RAT robust levelled off in the groundwater data, indicating robustness towards falsely detected taxa of the RAT robust workflow. With a minimum relative abundance cut-off of 0.001%, all rarefaction curves levelled off, although the different tools predicted different taxa richness. Kaiju estimated a much higher richness than RAT in both robust and sensitive mode (**Figure 4C**). Combined with the RAT results on simulated data where RAT robust underestimated richness while RAT sensitive included some false positives (**Supplementary Figure S6**), this shows that: (i) RAT robust is the best-suited RAT workflow in experiments where reliability is crucial, but it will likely not detect all taxa present while (ii) RAT sensitive will detect more taxa at the risk of including a few of them spuriously.

RAT estimates sequence abundance as opposed to taxonomic abundance [48]. This means that RAT reports the abundance of a taxon as fraction of total DNA in the sample, rather than as genome copies which can for example be estimated by querying marker genes [6–8]. The resulting relative abundance profile is skewed towards microorganisms with larger genomes, since they provide more DNA to the sequencing machine and thus contribute more reads than organisms with small genomes. To convert sequence abundance to genome copies, relative abundances have to be normalized by genome length, which is often unknown and can vary widely even between strains of the same species [49]. For novel microorganisms, genome sizes of closely related species might not be available. For these reasons, RAT by default does not convert sequence abundance to taxonomic abundance. However, the CAT pack provides a table with weighted mean genome sizes for most known bacteria and archaea at all taxonomic ranks based on genomes deposited in the BV-BRC database [50] which allows users to do this conversion if they wish.

## Conclusion

In this study, we presented the Read Annotation Tool (RAT), a new tool to strengthen the CAT pack metagenome analysis suite. We showed how annotating each read by using the best available taxonomic information (based on MAGs, contigs, or direct read mapping) leads to fewer falsely detected taxa and improves accuracy of taxonomic profiles. RAT is flexible to future improvements in sequencing technologies, as well as in assembly and binning software, as they are run by the user before the mapping and classification steps. RAT will be useful in the exploration and understanding of metagenomic datasets by robust classification of a majority of sequencing reads, even in unexplored environments that are rich in novel microorganisms.

## Methods

### RAT workflow

Read Annotation Tool (RAT) provides individual metagenomic sequencing reads with the most reliable taxonomic annotations, and uses these results to reconstruct an accurate taxonomic profile of the microbiome. RAT requires an input of sequencing reads, *de novo* assembled contigs or scaffolds, and optionally affiliated MAGs. We advise to filter the MAGs based on quality and only supply MAGs that have low contamination (<10%). Completeness of MAGs is less critical, as multiple contigs of the same organism carry a stronger taxonomic signal than individual contigs even if a part of the genome is not binned. These different DNA sequences are queried against a protein database for taxonomic annotations. Next, taxonomic annotations of individual reads are based on the associated data type with the highest confidence of annotation (MAGs > unbinned contigs > unassembled reads). RAT can be run in two different modes: sensitive (complete workflow, see below), and robust (skips step 3, only evidence from MAGs or contigs is used). The complete workflow of RAT consists of five steps:

1. RAT maps the reads back to the assembled contigs using bwa mem [51]. Reads mapping to each contig are extracted with samtools [52], including only primary mappings and excluding low-quality primary mappings (default: phred-score of 2, which can be changed by the user). In case of multiple mappings with equal phred scores, one of the mappings is assigned at random.
2. RAT performs taxonomic annotation of the contigs and MAGs with the previously published tools CAT and BAT [29], respectively. CAT and BAT annotate contigs and MAGs by predicting open reading frames (ORFs) with prodigal [32] and comparing these with DIAMOND blastp to the non-redundant protein database of NCBI (nr) [35] or the non-redundant set of proteins in GTDB [53], both of which can be downloaded and prepared by running ‘CAT prepare’. MAGs consist of binned contigs and therefore a contig in a MAG gets assigned both a BAT and a CAT annotation that may not be identical. As a MAG contains more taxonomic signals than a contig, RAT will prioritize the MAG annotation. In most metagenomic datasets, not all contigs are binned, and not all contigs can be annotated with CAT [22]. By default, RAT runs CAT with standard settings, and BAT with an f-value of 0.5 to prevent multiple annotations per MAG (see [22] for details). Currently, f-values < 0.5 are not supported by RAT.
3. Contigs that are not classified and reads that could not be mapped to any contig in step 1 are now classified simultaneously by comparing them to the protein database using DIAMOND blastx [33], and assigning the taxon of the last common ancestor of the organisms found within a certain range of the top hit (default: hits within 10% of the top-hit bit score, which can be changed by the user), similar to the -r parameter in CAT [22]. Thus, these direct mappings do not involve ORF predictions as in step 2.
4. Each individual read is classified according to the taxonomic signal with the highest confidence, in the following order: (i) If the read is mapped to a contig that is binned, the MAG annotation is assigned to it. (ii) If the read is mapped to an unbinned contig, the contig annotation is assigned to it. (iii) If a read is mapped to an unbinned contig that could not be annotated with CAT, or not mapped at all, the direct annotation is assigned to it (see step 3). (iv) Reads that do not have any taxonomic annotation are binned in an ‘unclassified’ category.
5. RAT calculates the abundance of a taxon by summing the total number of reads assigned to it, and normalizes abundances by dividing by the total number of sequenced reads in the sample. This final table constitutes the taxonomic profile. The relative abundances are sequence abundance (fraction of sequenced DNA), as opposed to taxonomic abundance (genome copies) [48]. A user may convert fraction of sequenced DNA to an estimate of genome copies by normalizing by genome size. The CAT pack provides a table with weighted mean genome sizes for most known bacteria and archaea at all taxonomic ranks based on a genomes deposited in the BV-BRC (previously PATRIC) database [50] which allows a user to do this conversion.

RAT is written in Python 3.8.3 [54] and available on Github at: https://github.com/MGXlab/CAT. We have tested RAT in the following configuration: bwa v. 0.7.17-r1188, samtools v. 1.10, prodigal v2.6.3, DIAMOND v. 2.0.5.

### Benchmarking on simulated datasets

To evaluate RAT’s performance as read classifier and taxonomic profiler, we used datasets generated for the second Critical Assessment of Metagenome Interpretation (CAMI) challenge [37]. We used 10 randomly selected samples of the mouse gut benchmark dataset (samples 6, 13, 23, 25, 26, 30, 33, 34, 38, and 53), which contain between 97-225 bacterial species each. We taxonomically annotated the reads with RAT and four other commonly used profilers: Bracken, Centrifuge, Kraken2, and Kaiju. All tools included in this benchmark also report relative abundance as sequence abundance and a comparison to RAT is thus fair [48].

For each read, we assessed at six taxonomic ranks (phylum, class, order, family, genus, and species) whether it was correctly or incorrectly annotated, or unclassified. To evaluate the taxonomic profiles, we used the same measures used in the original CAMI challenge [21]: the L1 and weighted Unifrac distances between the true and inferred profiles, as well as the precision and sensitivity of detected taxa. We only counted taxa as “detected” if they had been assigned at least 0.001% of the reads in the taxonomic profile and applied the same cut-off for all tools. L1 and weighted Unifrac are pairwise similarity measures between taxonomic profiles. L1 ranges from 0 (profiles are identical) to 2 (profiles do not share any taxa) according to the formula:

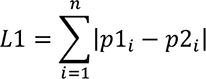

where *i* is the *i*th out of *n* total taxa in the union of the two profiles, and p1_i_ and p2_i_ are its relative abundances in the profiles that are being compared [21]. L1 is calculated at each taxonomic rank, contrary to weighted Unifrac distance. The weighted Unifrac distance incorporates both the relative phylogenetic or taxonomic relatedness between taxa and their abundance. We calculated weighted Unifrac distances using EMDUnifrac [55] using the taxonomy as measure for relatedness with a distance of 1 between taxonomic ranks as in [19]. Precision and sensitivity are defined as in [19] and only depend on the binary detection of each organism and not on their abundance. They were calculated using the following formulas:

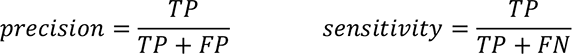

where TP (true positives) is the number of taxa that are correctly detected, FP (false positives) is the number of taxa that are incorrectly detected, and FN (false negatives) is the number of taxa that are not detected but are present in the dataset and thus should have been detected.

We ran Bracken v2.6.1, Kraken2 v2.1.2 [11], Centrifuge v1.0.4 [12], and Kaiju v1.8.2 [13] using default settings. We ran all tools using the nr/nt database from 08^th^ January 2019 that were provided by the CAMI challenge.

### Rarefaction curves

Rarefaction curves were calculated for RAT --sensitive, RAT --robust, and Kaiju results. We randomly sampled 1, 5, 10, 20, 30, 40, 50, 60, 70, 80, 90, and 100% of all reads ten times and counted the number of taxa detected in these subsets.

### Biological datasets

To demonstrate the performance of the RAT workflow on real-world data, we sequenced metagenomes from groundwater, a relatively unexplored biome [56], 18 samples were collected from three groundwater monitoring wells in an agricultural area in the Netherlands. The same samples were used in an earlier study where they were analyzed with 16S rRNA amplicon sequencing [38]. Each well was sampled at six discrete depths between 12 and 64m below the surface. We filtered 5-7L of groundwater through 0.2µm filters and extracted DNA from the filters using the DNeasy PowerSoil Kit (Qiagen, Hilden, Germany) according to the manufacturer’s instructions. DNA quality (average molecular size) was checked with 1% (w/v) agarose gels stained with 1x SYBR® Safe (Invitrogen, Grand Island, NY) and quantified using the dsDNA HS Assay kit for Qubit fluorometer (Invitrogen). Whole metagenome shotgun sequencing was performed on the DNA by Novogene in Hong Kong on the Illumina MiSeq Platform, generating 35,289,790-58,902,006 bp paired-end sequencing reads per sample (**Supplementary Table ST1**).

For quality-control, assembly, and binning, we used the ATLAS pipeline v2.4 [57]. ATLAS uses BBTools [44] to remove PCR duplicates and adapters and to trim the reads, assembles the reads using SPAdes v3.13.1 in metagenomic mode [25], and bins the contigs using MetaBAT2 v2.14 [27] and maxbin2 v2.2.7 [28], after which DASTool v.1.1.2 [45] is used to optimize MAGs resulting from the two binning approaches. We used MAGs of medium- to high-quality (>50% completeness, <10% contamination [46]) based on CheckM estimates in lineage-wf mode [43]. We ran RAT on multiple samples at a time using GNU parallel v20210622 [58] with the nr database from the 04^th^ of March 2020. We ran Kaiju on the reads using default settings with a database containing NCBI nr for bacteria, archaea, viruses, fungi and microbial eukaryotes from the 24^th^ of February 2021.

The N50 of the assembled contigs was between 1435-3012nt per sample, the L50 was between 15,196-39,823nt. Out of the 2,770,251 total contigs that were generated from the 18 samples, CAT annotated 2,411,810. All 423 medium- to high-quality MAGs were annotated at superkingdom rank or lower by BAT.

To further assess the diversity of groundwater organisms represented by the MAGs, we dereplicated all medium- to high-quality MAGs with dRep using default settings [47]. We performed a phylogenetic analysis of the dereplicated MAGs based on the CheckM alignment of 43 universal marker genes that are used for phylogenetic placement [43]. A phylogenetic tree was inferred with IQ-TREE v2.1.2 [59], ModelFinder [60] and UFBoot [61], using the model LG+R10 chosen according to BIC. The resulting tree was visualized with iTOL [62]. The tree was rooted between the archaeal and bacterial MAGs based on their BAT classification (by RAT).

### Plotting

All figures were made using R 4.1.3 and Rstudio v. 1.1.456 [63]. The packages used for plotting were ggplot2 [64], tidyverse [65], reshape2 [66], ggalluvial [67], dplyr [65], tidyr [68], RColorBrewer [69], Hmisc [70], vegan [71], ape [72], and gridExtra [73].

## Declarations

### Ethics approval and consent to participate

Not applicable.

### Consent for publication

Not applicable.

### Availability of data and materials

RAT is available on github under https://github.com/MGXlab/CAT. The scripts used in the downstream analyses are available on github under https://github.com/thauptfeld/RAT_paper. The raw sequencing reads used for the biological analysis are available at SRA under BioProject ID PRJNA947390. Data from the CAMI challenge is available at https://data.cami-challenge.org/participate.

### Authors’ contributions

EH developed RAT, integrated it into the CAT_pack, performed the CAMI benchmarks, and wrote the manuscript. NP integrated the GTDB option into CAT prepare. SvI and BLS performed the analysis on biological data. AAV sampled the groundwater, extracted DNA and sent the groundwater samples for sequencing. BED and FABvM supervised the research and co-wrote the manuscript.

## Acknowledgements

We thank Nora B. Sutton and her group for providing us with groundwater metagenomes to use for this manuscript. We thank Jan Kees van Amerongen for critical technical support. We thank the members of TBB at the University of Utrecht for their valuable input on the text.

---

**Supplementary Figure S1.**
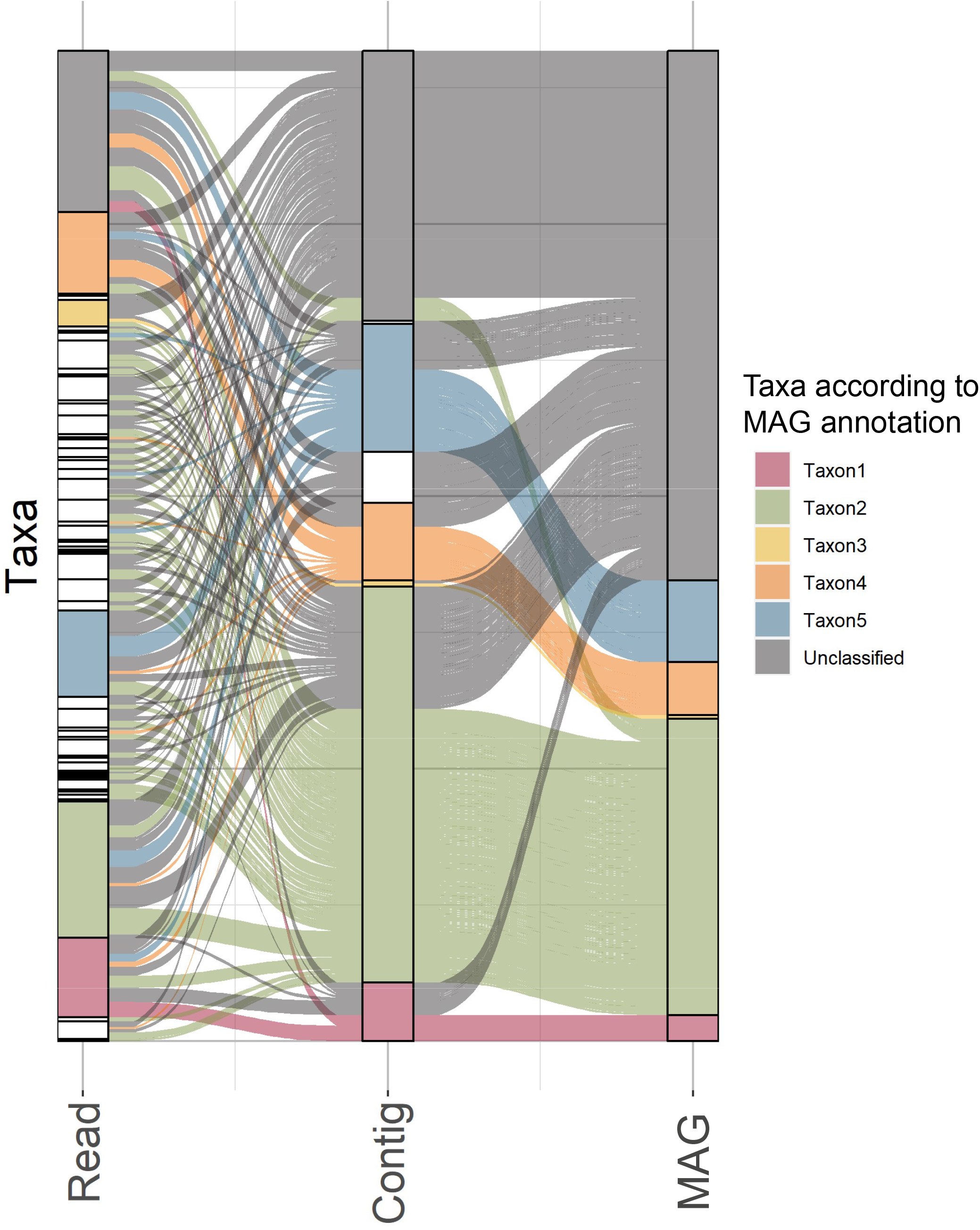
Schematic depiction of noise reduction by using reliable taxonomic signals. Each column segment represents a taxon, each column represents an annotation step. In the read annotation step, many taxa are detected, the profile is noisy. At contig level, the number of detected taxa is much lower. Reads that were previously unannotated or had a spurious annotation now get annotated to one of eight main taxa. However, many reads that had an annotation on read level do not get an annotation on contig level, either because they don’t map to any contigs, or because they map to a contig without annotation. At MAG level, there are even fewer detected taxa, and more unclassified data.

**Supplementary Figure S2.**
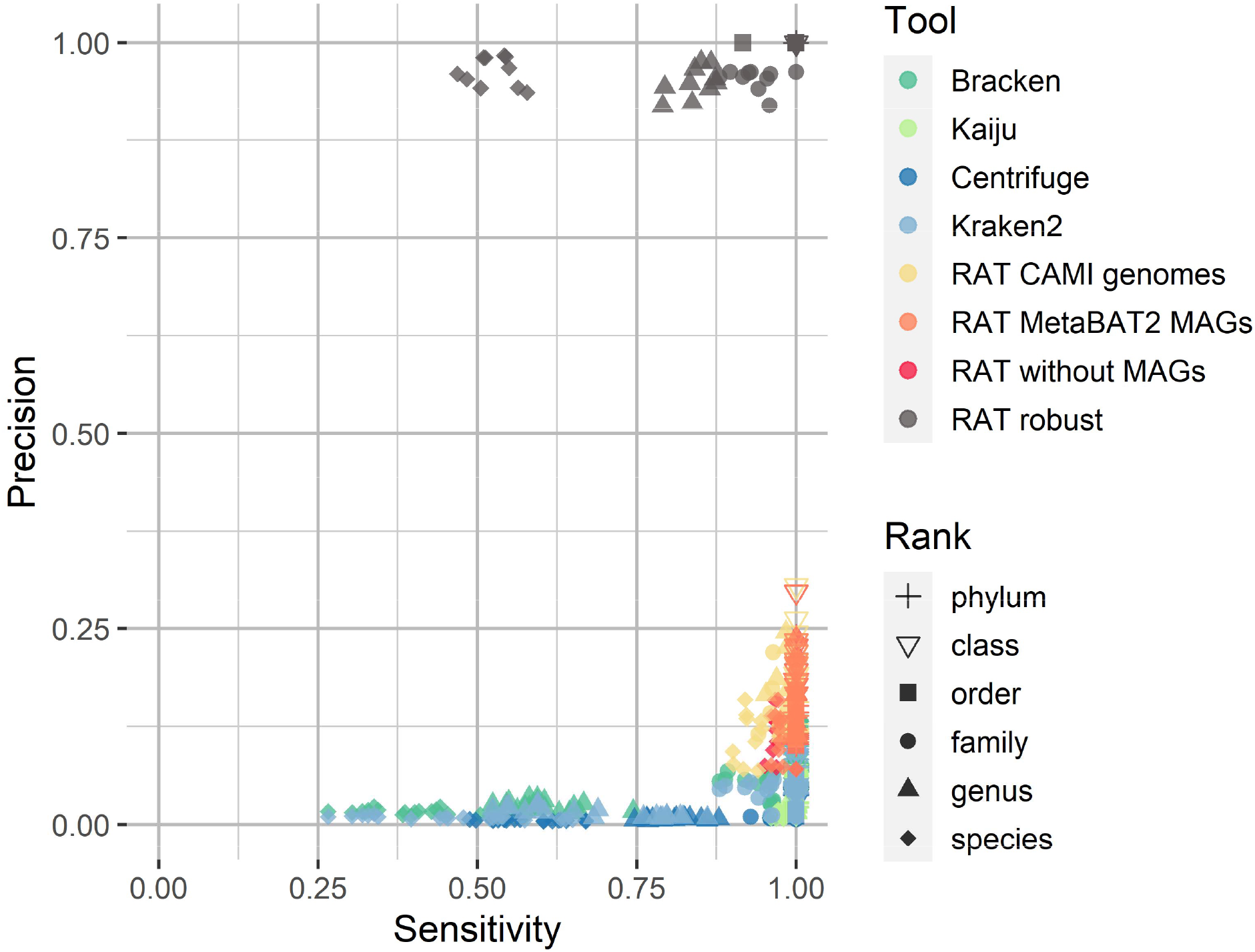
Sensitivity vs. Precision of the different tools without an abundance cut-off to include taxa. Different shapes signify the sensitivity/precision on different ranks, different colors indicate tools (high precision + sensitivity is better).

**Supplementary Figure S3.**
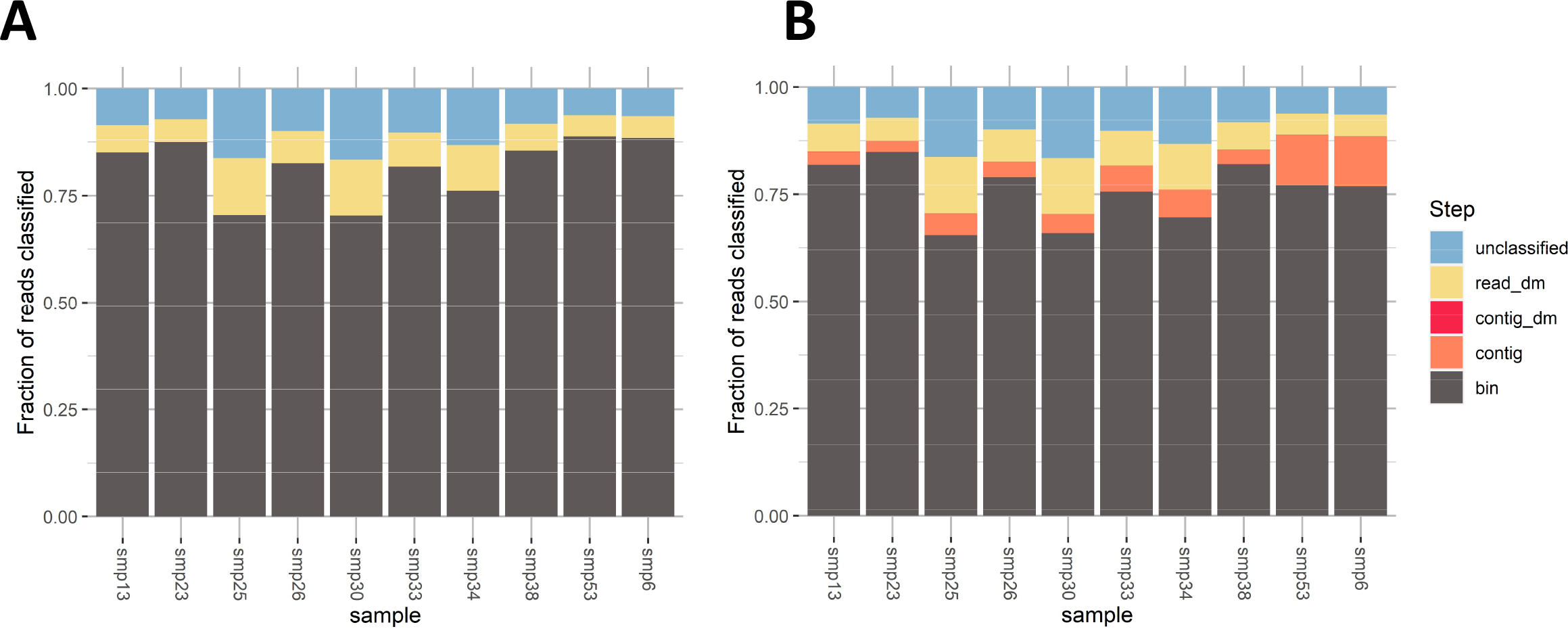
Fraction of read annotations in the simulated CAMI dataset and the taxonomic signal they originate from. ‘bin’ refers to BAT annotation of a MAG, ‘contig’ refers to CAT annotation of a contig, ‘contig_dm’ and ‘read_dm’ refer to diamond blastx direct annotations of contigs/reads. (A) Read annotations per taxonomic signal using CAMI genomes as MAG input. (B) Read annotations per taxonomic signal using MAGs binned by MetaBAT2.

**Supplementary Figure S4.**
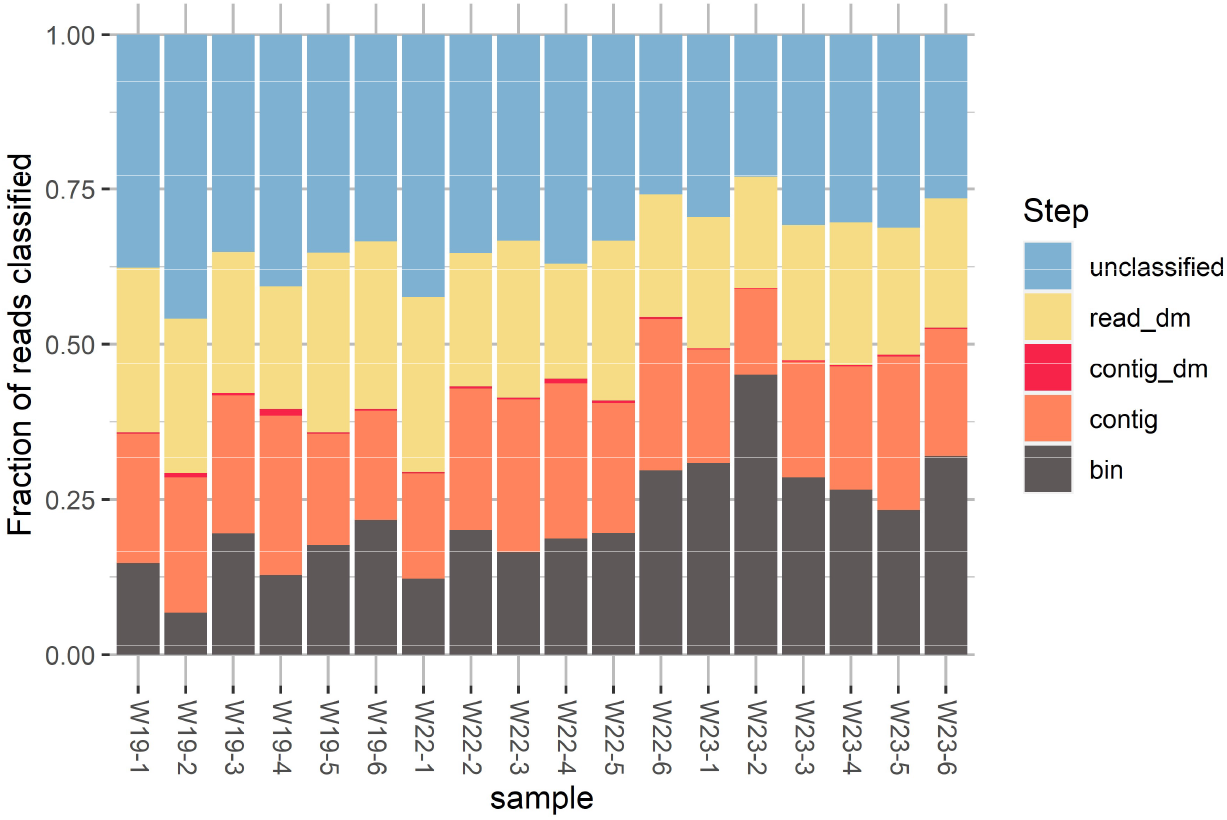
Fraction of read annotations in the biological groundwater dataset and the taxonomic signal they originate from. ‘bin’ refers to BAT annotation of a MAG, ‘contig’ refers to CAT annotation of a contig, ‘contig_dm’ and ‘read_dm’ refer to diamond blastx direct annotations of contigs/reads.

**Supplementary Figure S5.**
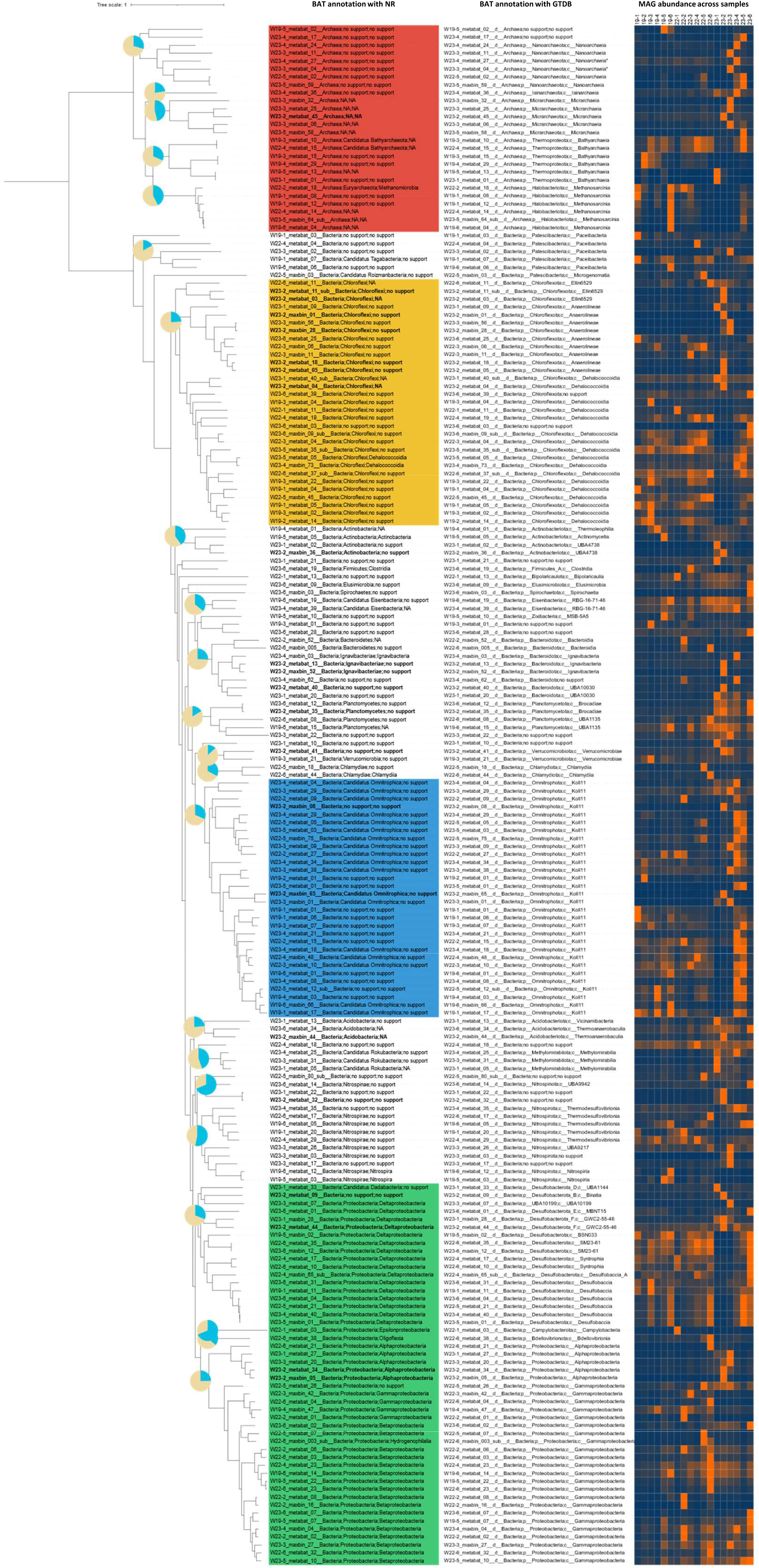
Phylogenetic tree of MAGs assembled from groundwater samples based on checkM universal marker genes. MAGs were dereplicated with dRep. The pie charts indicate fraction of reads per phylum that are mapped to MAGs (in blue) or not mapped to MAGs (in beige). MAG annotations are on superkingdom, phylum and class level, based on BAT annotation with NR (left) and GTDB (right). The heatmap on the right indicates how abundant the MAG is in each sample (normalized, max per row is 1/orange). Abundance is based on reads that map to each MAG or another MAG in the same dRep cluster per sample.

**Supplementary Figure S6.**
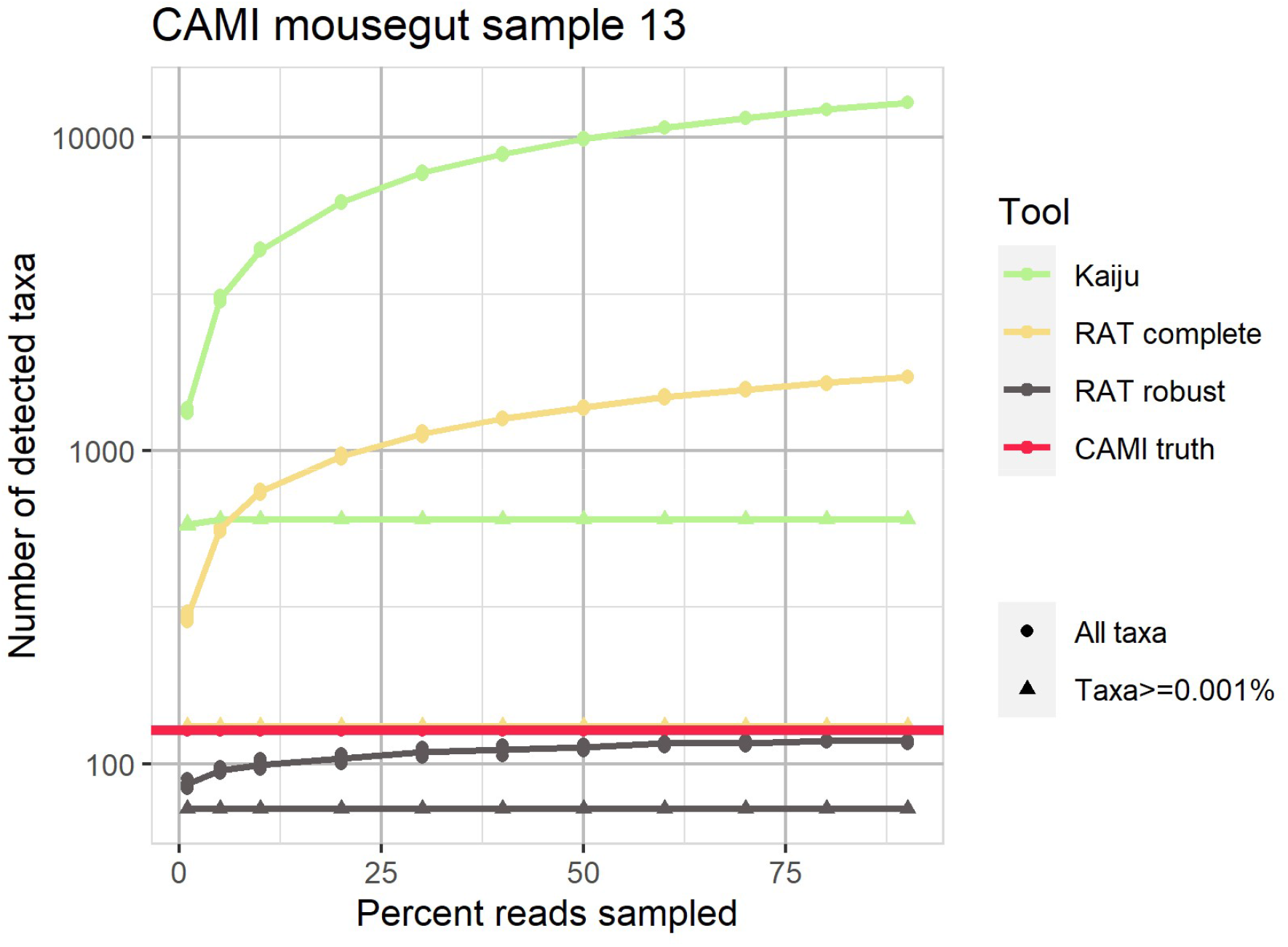
Number of taxa detected in one of the simulated CAMI datasets by RAT robust, RAT sensitive, and Kaiju. ‘CAMI truth’ refers to the actual number of taxa present in the sample. Triangles indicate the number of taxa detected in profiles when a minimum abundance is required to consider an organism as detected. Circles indicate the number of taxa detected without a cut-off.

